# Functional Extinction of a Genus of Canopy-Forming Macroalgae (*Cystophora* Spp.) Across its Warm Range Edge

**DOI:** 10.1101/2022.09.08.507225

**Authors:** Albert Pessarrodona

## Abstract

Declines of canopy-forming algae are increasingly prevalent in temperate latitudes, although most losses have occurred across densely populated areas. Here, I use a combination of field surveys, anecdotal evidence, records from the literature and herbaria collections, to document the biogeography of the canopy-forming genus *Cystophora* across the sparsely populated end of its distribution in Western Australia. Although historically common, most species were found to be absent or exceedingly rare across their rear (warm) range edge, suggesting their functional extinction. Out of five species, three experienced range contractions >500 km, with some losing approximately 8% of their global distribution. Reasons for their decline are unknown, but likely involve gradual warming, marine heatwaves and rapid urbanization. Increasing human impacts and further warming in the region threatens several species with further extirpation, some of which are endemic to the area and play unique ecological roles.

## Main text

Subtidal canopy-forming kelps and fucoids form impressive underwater forests across temperate and polar latitudes. Declines and local extirpations of these seaweeds are however becoming increasingly prevalent as anthropogenic impacts increase on the marine environment (Thibaut *et al*., 2005; Krumhansl *et al*., 2016; McPherson *et al*., 2021), with negative consequences for ecosystem functioning and their associated fauna and flora. Losses and local extirpations have been particularly widespread in areas with intense human pressure such as heavily urbanized coasts (Kautsky *et al*., 1992; Thibaut *et al*., 2005), with non-urbanized areas avoiding most declines (Benedetti-Cecchi *et al*., 2001; Scherner *et al*., 2013). As the coastal development and urbanization rates rapidly increase across the worlds coastlines (Seto *et al*., 2011), there is an urgent need to survey canopy-forming algae biodiversity in less populated areas. Yet, studies on subtidal canopy-forming algae in those areas have mostly focused on charismatic and well-studied species (mostly kelps), with little being known about other canopy-forming groups (Shepherd & Edgar, 2013).

Southern Australia features one of the richest marine floras in the world, including an unparallel and highly endemic diversity of canopy-forming algae (>60 species). A range of anthropogenic pressures have precipitated declines and local extirpations of several of these species (Coleman *et al*., 2008; Connell *et al*., 2008; Shepherd *et al*., 2009; Phillips & Blackshaw, 2011; Smale & Wernberg, 2013; Wernberg *et al*., 2016), with ocean warming threatening severe range contractions in the coming decades (Martínez *et al*., 2018). Of particular concern is the predicted contractions of species in the genus *Cystophora*, which is endemic of Australasia, extremely diversified (23 species), and play unique functional roles distinct from other canopy-forming seaweeds (Coleman & Wernberg, 2017). *Cystophora* plants often occur as mixed stands with no single species dominating the assemblage (Turner & Cheshire, 2003), much like trees in tropical rainforests. Despite being the prevalent canopy-forming algae along much of the southern Australian continent (Turner & Cheshire, 2003; Goldberg & Kendrick, 2004), little is known about their ecology and fine-scale biogeographic distribution. Here, I combine presence/absence records from herbaria and field surveys to show that the entire genus has largely disappeared from the warmer range of its distribution in Western Australia.

To establish the historical biogeographical distribution of *Cystophora* across its western range edge, I conducted a comprehensive search of *Cystophora* records in herbaria, government reports and peer-reviewed studies. Historical herbaria collections were sourced from the Australasian Virtual Herbarium (AVH, 2021) and the Macroalgal Herbarium Consortium Portal (macroalgae.org), which contains digitized information of herbaria stored in the United States. I also searched for records in the Natural History Museum (UK) and the Muséum National d’Historie Naturelle (France) given that many French and British botanists collected macroalgal specimens from Australian coasts (Womersley, 1959). However, specimens in those collections were not properly georeferenced and therefore not included. More recent presence/absence records were gathered from the grey literature (Gordon, 1986; Moore, 1987; Burt & Anderton, 1997; Colman, 1997; Bancroft & Davidson, 1998; Kendrick *et al*., 1999; Westera *et al*., 2007; Edgar *et al*., 2009), published datasets (CSIRO, 2005), peer-reviewed studies (Kendrick *et al*., 2004; Waddington *et al*., 2010; Richards *et al*., 2016), and unpublished theses (Smith, 1952; Wood, 1980; Phillips, 1996; Scott, 2012). Finally, a series of 30 min underwater visual censuses targeting *Cystophora* along the entirety of the coast— including revisits of many of the historical collection sites— were conducted between 2018 and 2021 to determine the more recent extent of *Cystophora* species across the entirety of their range (57 reefs from Geraldton to Esperance).

According to historical reports and specimens deposited in herbaria (Lucas, 1936; Womersley, 1964, 1987; AVH, 2021), a total of fourteen *Cystophora* species occur in Western Australia, with six species (*C. brownii*, *C. grevillei* and *C. monolifera*, *C. pectinata*, *C. subfarcinata*, *C. retroflexa*) present in the warm range edge (i.e. >20°C annual mean surface temperature, ca. above 32.5°S latitude; Fig. 1). Of these, *C. brownii*, *C. grevillei* and *C. monolifera* extended as far north as Port Denison/Geraldton (29°S 115°E), many of those specimens being collected or identified by the eminent Australian phycologist H.B.S Womersley. The pioneering studies around the Perth Metropolitan area (32°S 115°E) by G. G. Smith (1952) described five species (all but *C. subfarcinata*) being present as “typical subordinate species of the [subtidal] reef system” and “very commonly found as drift” during the 1940s, with *C. monilifera* and *C. brownii* also extending to the intertidal-subtidal fringe. During the 1950s-1970s, *Cystophora* plants were frequently collected (>100 records) as drift by phycologists at a variety of locations across the warm range edge (e.g. Dongara, 29°S 115°E; Yanchep, 31°S 115°E; Fremantle, 32°S 115°E), as well as throughout the colder South coast (>300 records; Fig. 1) (Hodgkin, 1959a,b; Womersley, 1964).

**Fig. 1.**
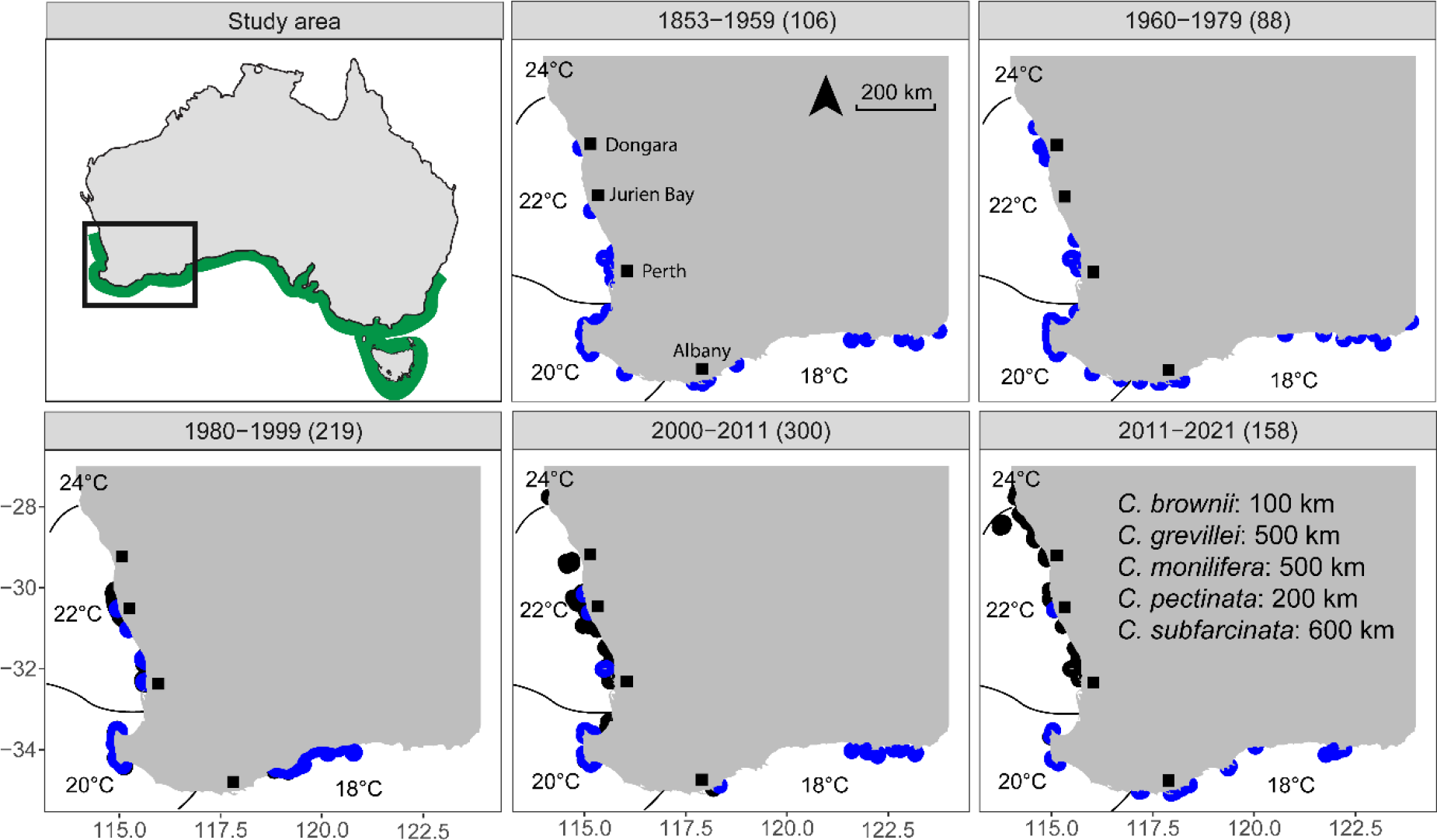
Occurrence of *Cystophora* spp. across the western range edge of its distribution (depicted in green) over time. Presences are indicated in blue, while black denotes absences in surveyed locations. The number of sampling events (i.e. a survey at a site at a given year) is indicated in parenthesis next to the different study periods. Lines show the average annual mean sea surface temperature isotherms (1961-1990). The extent of putative range contractions of several *Cystophora* spp. are shown in the final panel. Note that pre-1980s records are mostly from herbaria collections and therefore only presences are reported.

By the time some of the first subtidal work on SCUBA was conducted in the 1980s, the number of records in the warm range edge had drastically declined (4 records across 136 sampling events at 136 sites, Fig. 2), when some of the first subtidal work on scuba was conducted. Intense sampling (40 sites) of the algal communities in the Marmion lagoon (32°S 115°E) for the establishment of the state’s first marine park failed to detect any species (Moore, 1987), and so did studies conducted at Perth’s nearshore reefs similar to the ones where they were historically frequent (Phillips, 1996). The early 2000s saw a sharp increase in the algal survey efforts which translated in an increase in *Cystophora* records across the entire coast (>300 records across 300 sampling events at 228 sites; Fig. 2), but not in the warm range edge (7 records across 191 sampling events and 147 sites). A few sparse individuals of *C. brownii* (0.8-1.7% cover) were recorded at two of the 42 sites surveyed between Jurien Bay and Lancelin (30-31°S 115°E; Barrett *et al*., 2002; Edgar *et al*., 2005), and single specimens of *C. grevillei* and *C. monilifera* were recorded in Rottnest Island (32°S 115°E; Scott, 2012; AVH, 2021). It thus appears that *Cystophora* was rare or had disappeared from the most of its warm range edge by the early 2000s.

**Fig. 2.**
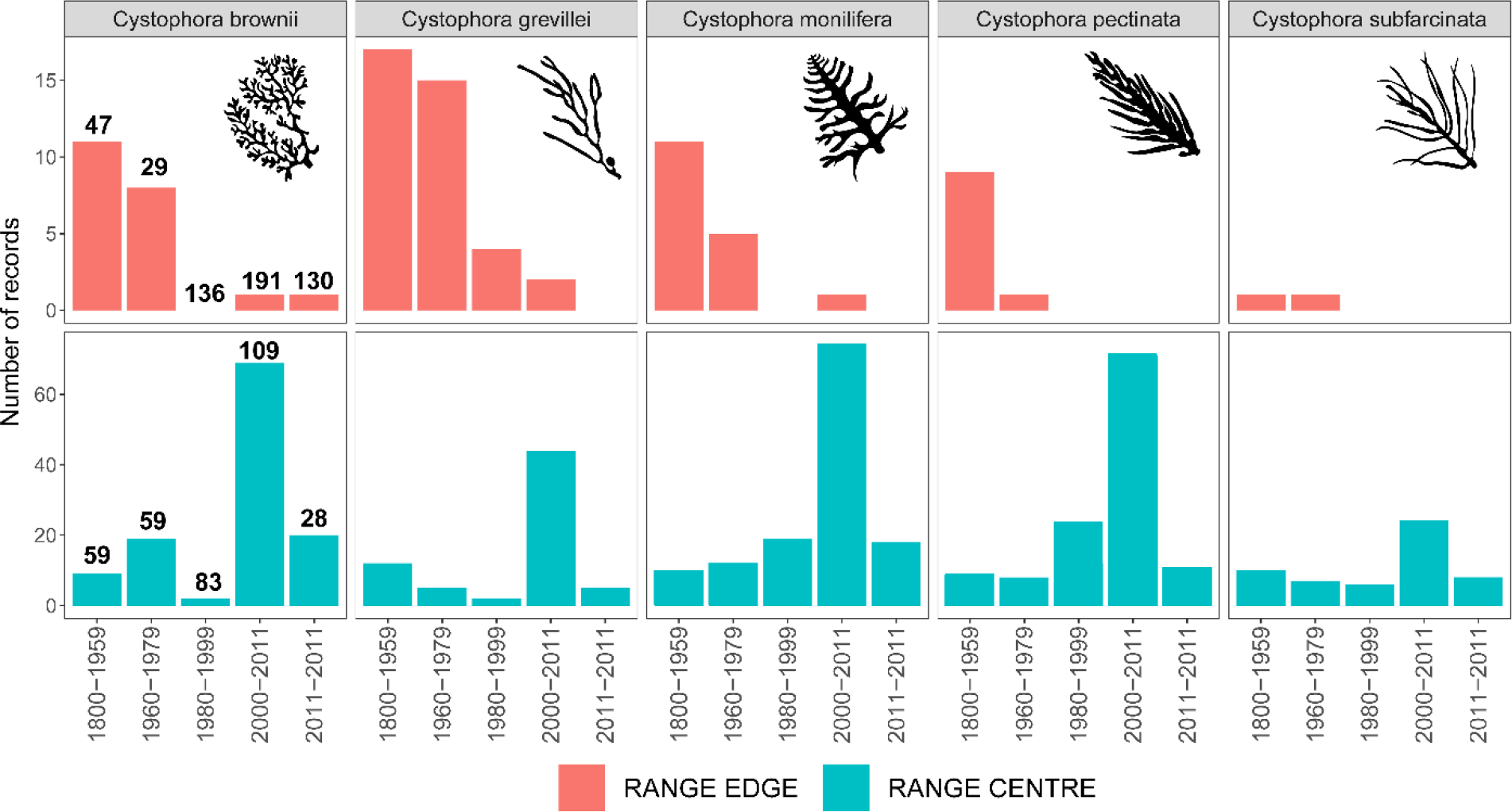
Number of records of *Cystophora* spp. over time along the western rear (warm) edge (>20°C annual mean surface temperature, ca. above 32.5° S latitude) and central (cool) range (<20°C, below 32.5° S latitude). Records include herbaria collections, field algal surveys from the peer-reviewed and grey literature, as well as underwater visual censuses targeting *Cystophora* across the entirety of its range (2018-2021). The number of sampling events in the range edge and centre during each time period is indicated in bold on top of the bars, and is shared by each species.

The 2010s were marked by a prolonged marine heatwave that negatively affected Western Australian marine ecosystems, driving the range contraction of several cold-water seaweeds (Smale & Wernberg, 2013; Wernberg *et al*., 2016). A comprehensive set of algal surveys along 52 reefs conducted immediately after the heatwave only recorded *Cystophora* at the cooler (i.e.<20°C annual mean surface temperature) portions of its distribution (Bennett, 2015). During the *Cystophora* visual censuses I conducted a decade after the heatwave (2018-2021), *Cystophora* was absent from many of the historical collection sites in the warm range edge (Port Denison, 29°S 115°E; Mudurup Reef, Cottesloe, 32°S 115°E; Point Peron, 32°S 115°E), being present at only one of the 28 reefs that were surveyed in the region. In contrast, all *Cystophora* species occurring in Western Australia were recorded from the cooler distributional range. Although it remains possible that small, isolated populations exist in warm areas that were not sampled, as suggested by the few isolated *C. brownii* plants (ca. 1 plant m^−2^) that the author recorded in one of the nearshore dives conducted in Cervantes (30.5°S 115°E), it appears that *Cystophora* species may be functionally extinct across ca. 100-600 km of their historical range edge. In the case of *C. pectinata*, this contraction equates to a loss of approximately 8% of its global distribution.

The approach used here necessitated collection of records from several sources, which may have varied in sampling effort. Herbarium records rarely represent systematic collecting efforts, and so it is not possible to estimate the historical *Cystophora* abundance throughout their warmer range nor determine whether they had a continuous distribution. The frequency in which they were encountered along the coast however suggests that they were common, at least until the past half-century (Smith, 1952; Hodgkin, 1959a). An additional issue with collating presence/absence data from surveys with multiple objectives is that untrained scientists may have failed to identify *Cystophora* (i.e. false absences), yielding misleading impressions of their decline. This is however, highly unlikely. Many of the surveys conducted during that time exclusively targeted algal assemblages (Phillips, 1996; Wernberg *et al*., 2003; Westera *et al*., 2007; Smale *et al*., 2010; Bennett, 2015), some of which recorded upwards of 250 species of macroalgae. The sampling effort across the warm range edge also increased massively in terms of spatial cover and number of sampling events (>400 sampling events at >300 sites after the 1980s versus 76 sampling events at 71 sites before). Additionally, surveys conducted at the same area on multiple occasions reported *Cystophora* repeatedly, even when algae was not the sole focus on the survey (Edgar *et al*., 2009). Finally, the visual censuses that I conducted specifically targeted *Cystophora* and included some of the historical collection sites, reinforcing evidence that the genus has declined substantially.

While declines and local extirpations of canopy-forming macroalgae have been recorded worldwide (e.g. Coleman *et al*., 2008; Connell *et al*., 2008; Phillips & Blackshaw, 2011; Thibaut *et al*., 2015), the apparent range retractions documented here are particularly unique in that they involve the functional extinction of multiple species within the same genus, and that some occurred across relatively large sections of unurbanized coast (e.g. outside Perth Metropolitan area). While it is not possible to determine the causes of their demise, there are several non-mutually exclusive possibilities. Summer temperatures are the best predictor of *Cystophora* spp. biogeographical distributions (Martínez *et al*., 2018), suggesting that they are highly sensitive to warming (Bennett, 2015). Since the 1950s, the waters around the Western Australian continental shelf have warmed about 0.9°C (Pearce & Feng, 2007) and experienced an extreme marine heatwave in 2011 that affected large portions of the coastline and precipitated multiple seaweed range contractions (Smale & Wernberg, 2013; Wernberg *et al*., 2016). It thus seems plausible that warming is one of the lead causes of their decline. Another possible driver of decline is decreasing water quality associated with urbanization. While the majority of the Western Australian coast remains relatively unurbanized, development since the 1950s has concentrated in the Perth metropolitan area, whose population has grown sevenfold. Multiple ocean outlets have been added since the 1960s to satisfy the increasing wastewater supply, which discharged ca. 80 Glitres year^−1^ around the 2000s (Water Corporation). While the tolerance of *Cystophora* to increased nutrients is unknown, other Australian subtidal fucoids are highly sensitive to treated sewage effluents (Brown *et al*., 1990). Along sheltered areas and embayments, increases in sedimentation have driven declines in some *Cystophora* spp. (Shepherd *et al*., 2009). This is unlikely to be the main driver in this scenario however, as reefs in the region are exposed to intense wave action and large oceanic swells (Hemer, 2006), which generally maintain low levels of turbidity and sedimentation.

Recovery of *Cystophora* spp. across their former range seems unlikely given that only reduced populations of few individuals like the one encountered in Cervantes possibly remain, severely restricting the available propagule supply. The large distances between remaining populations may further prevent recolonization and connectivity. Over 90% of the propagules from fucoids settle within metres of the parental thalli (Kendrick & Walker, 1995), *Cystophora* recruitment being negligible 500 m away from reproductive plants (Goldberg *et al*., 2004). The functional extinction of *Cystophora* along its warm range edge is concerning, as they play functionally unique roles in the ecology of temperate reefs (Coleman & Wernberg, 2017), supporting different faunal assemblages and having different growth than the more well-studied kelp species for example (A. Pessarrodona & C. Grimaldi unpublished data). Further warming and coastal development threatens populations established along the cooler southern coast (Martínez *et al*., 2018), which is concerning given that several species are endemic to that area (e.g. *C. gracilis*,*C. harveyi*), and have extremely narrow geographical distributions (e.g. 300 km). A better understanding of their ecology is key to predict their response to increasing anthropogenic pressures and prevent further declines.

## Acknowledgments

The author declares no competing interests

